# Novel diversity within marine Mamiellophyceae (Chlorophyta) unveiled by metabarcoding

**DOI:** 10.1101/449298

**Authors:** Margot Tragin, Daniel Vaulot

## Abstract

The Ocean Sampling Day (OSD) project provided metabarcoding data for the V4 hyper-variable regions of the 18S rRNA gene from 157 samples collected at 143 mostly coastal stations. In this paper we focus on the class Mamiellophyceae, which was found at nearly all OSD stations and represented 55 % of the green microalgae (Chlorophyta) reads in the 2014 OSD dataset. We performed phylogenetic analyses of unique OSD metabarcodes (ASV, amplicon single variants) and reference GenBank sequences from cultures and from the environment, focusing on the four most represented genera: *Ostreococcus* (45 % of the Mamiellophyceae reads), *Micromonas* (34 %), *Bathycoccus* (10 %) and *Mantoniella* (8.7 %). These analyses uncovered novel diversity within each genus except *Bathycoccus*. In *Ostreococcus*, a new clade (E) with 2 very clear base pair differences compared to the oceanic clade B in the V4 region was the second most represented clade after the coastal *Ostreococcus* “*lucimarinus*”. Within *Micromonas*, ten clades were found exceeding the 4 species and 2 candidate species already described. Finally, we found 2 new environmental clades of *Mantoniella*. Each Mamiellophyceae clade had a specific distribution in the OSD dataset suggesting that they are adapted to different ecological niches.

## Introduction

In marine waters, the accepted paradigm is that the so-called “red” lineage (mainly Diatoms and Dinoflagellates) is dominant, while the “green” lineage (land plants) is dominating the terrestrial environments^1^. These two lineages differentiate by the evolutionary origin of their chloroplast: those of the “green” lineage are surrounded by two membranes, which is an evidence for a single endosymbiotic event and their major photosynthetic pigments are chlorophyll a and b^1^. Studies in coastal waters with both microscopic and molecular techniques found that the “green” lineage, which is mainly represented by Chlorophyta among unicellular protists, is dominant in these ecosystems^2–4^. Metabarcoding studies following the development of high throughput sequencing (HTS) have confirmed the importance of the Chlorophyta in marine waters^5,6^.

The European Ocean Sampling Day project (OSD) sampled the world coastal waters in 2014 around the summer solstice (21 June) with the aim of analyzing the diversity and distribution of marine microorganisms^7^ especially in coastal waters using 18S rRNA metabarcodes (V4 and V9 hyper-variable regions)^8^. This dataset allowed to determine the distribution of fourteen classes of Chlorophyta^9^. Mamiellophyceae^10^, the most prevalent class in all coastal environments^11^, did not show any geographic distribution patterns or environmental preference^9^. We hypothesize that, in order to detect distribution patterns, this class should be investigated at lower taxonomic levels, in particular the species level.

Mamiellophyceae consist of four orders: Mamiellales, Bathycoccales, Dolichomastigales and Monomastigales^10^. Monomastigales are confined to freshwater environments while Dolichomastigales, although quite diversified in marine waters^12,13^, are a minor component of Mamiellophyceae in coastal waters. In contrast, Mamiellales and Bathycoccales host some of the most common Chlorophyta microalgae such as the ubiquitous *Micromonas*, the smallest known eukaryote *Ostreococcus* or the *coccoid Bathycoccus*^10^. Within Mamiellales, *Micromonas pusilla*^14^ was recently split into four species: *Micromonas bravo* (previously clade B.E.3), *Micromonas commoda* (previous clade A.ABC.1-2), *Micromonas polaris* (previously clade B arctic), *Micromonas pusilla* (previously clade C.D.5) and two clades mentioned as candidate species 1 (clade B. .4) and candidate species 2 (clade B warm)^15^. Within the genus *Mantoniella*, only two species have been described: the ubiquitous *Mantoniella squamata*^3,16^, first described as *Micromonas squamata*^17^, and *Mantoniella antarctica*^18^. Within Bathycoccales, four *Ostreococcus* clades have been delineated^19^: *Ostreococcus tauri*^20^, *Ostreococcus mediterraneus*^21^, both of which have been formerly described, *Ostreococcus* “*lucimarinus*” (clade A) and clade B, which both lack formal taxonomic description. Analyses of pigment content and response to light levels allowed to distinguish two broad ecotypes: strains adapted to high (*O. tauri,O. mediterraneus, O. “lucimarinus”*) and low (*Ostreococcus* clade B) light^22^. The second genus within Bathycoccales hosts a single species, *Bathycoccus prasinos*^23^. No clades can be delineated inside this species based on 18S rRNA gene sequences from cultures and the environment. However, divergence in ITS sequences suggest that *Bathycoccus prasinos* probably consists of two different species^24,25^. The present paper uses the OSD V4 dataset to analyze the taxonomic diversity and global distribution of four major Mamiellophyceae genera: *Ostreococcus, Micromonas, Bathycoccus* and the less studied *Mantoniella*. Our analyses reveal the existence of novel clades within *Ostreococcus* and *Micromonas* and allowed to determine that most species/clades have specific oceanic distributions.

## Material and Methods

The Ocean Sampling Day consortium provided 2 metabarcoding datasets for 2014 using the V4 region of the 18S rRNA gene: the LGC dataset consisting of 157 water samples from 143 stations (see Table S1) filtered on 0.22 *µ*m pore size Sterivex without prefiltration and the Life Watch (LW) dataset consisting of a subset of 29 water samples filtered on 0.8 µm pore size polycarbonate membranes without pre-filtration. The extraction, PCR and sequencing protocols have been described previously^8,9^. In brief, the LGC and LW data originated from the same water samples but have been processed independently for filtration, DNA extraction, PCR amplification and sequencing (both Illumina 2×250 bp). LGC and LW provided respectively about 5 and 9 millions of V4 sequences, resulting in higher coverage for the LW dataset.

The LGC and LW datasets (https://github.com/MicroB3-IS/osd-analysis/wiki/Guide-to-OSD-2014-data) were analyzed with the same pipeline using the Mothur software v. 1.35.1^26^. Reads were filtered to keep only sequences without ambiguities (N) and longer than 300 pb. Reads were aligned on SILVA seed release 123 alignment^27^ corrected by hand to remove gaps at the beginning and at the end of alignments. The aligned datasets were filtered by removing columns containing only insertions. Chimeras were checked using Uchime v. 4.2.40^28^ as implemented in Mothur. Unique sequences (ASVs or Amplicon Single Variants^29^) were assigned using the Wang classifier as implemented in Mothur and the PR^2^ reference database^30^ version 4.2 (https://figshare.com/articles/PR2rRNAgenedatabase/3803709/2) for which the Chlorophyta sequences had been recently curated^6^.

To check the assignation and explore the genetic diversity of Mamiellophyceae, ASVs from *Bathycoccus, Micromonas, Ostreococcus* and *Mantoniella* represented by more than 200 reads for either LW and LGC were aligned to GenBank reference sequences using MAFFT v. 7.017^31^. Maximum likelihood (ML) phylogeny was built using Fast Tree v. 1.0^32,33^ as implemented in Geneious v. 7.1.9^34^. Bayesian phylogeny was built with Mr Bayes v. 3.2.6.^35^ implemented in Geneious. Clades were defined both with the presence of clear signatures in the alignments and by phylogenetic features^6,36,37^): monophyletic assemblages of sequences in the phylogenetic trees, well supported (> 70 % bootstrap), and found with the two different construction methods (i.e. maximum likelihood and Bayesian).

Relative abundance of selected ASVs was computed using R software version 3.3.1 (http://www.R-project.org/) on the LGC dataset for a subset of 92 samples with more than 100 Mamiellophyceae reads. Graphics were performed using the R packages ggplot2, ComplexHeatmap^38^ and treemapify.

## Results and Discussion

We analyzed unique sequences (ASVs for Amplicon Single Variants^29^) of two separate OSD data sets: LGC and LW. We focused on the LGC dataset which encompasses a much larger number of samples than the LW dataset which corresponds to a subset of the OSD samples that have been processed in a completely independent manner compared to the LGC (different filtration, DNA extraction, PCR and Illumina sequencing). The LW dataset was used mainly to confirm the fact that the LGC ASVs were not artefacts by verifying that to any major LGC ASV corresponded a LW ASV with a strictly identical sequence.

Mamiellophyceae represented 55 % of the Chlorophyta reads found in the OSD 2014 surface samples^9^. Seven described Mamiellophyceae genera were recovered among LGC ASVs while the other ASVs could only be classified at the family or order level. Four genera were clearly dominant: *Ostreococcus, Micromonas, Bathycoccus* and *Mantoniella*, the two former with almost equal contribution (Fig. 1). We decided to focus on the 23 major LGC ASVs for these four genera that were represented by at least than 200 reads (Table 1) corresponding 68% of the Mamiellophyceae reads. Among these ASVs, *B. prasinos* was clearly the most ubiquitous followed by 2 *Micromonas* clades and O. “*lucimarinus*” (Fig. S1).

**Figure 1.**
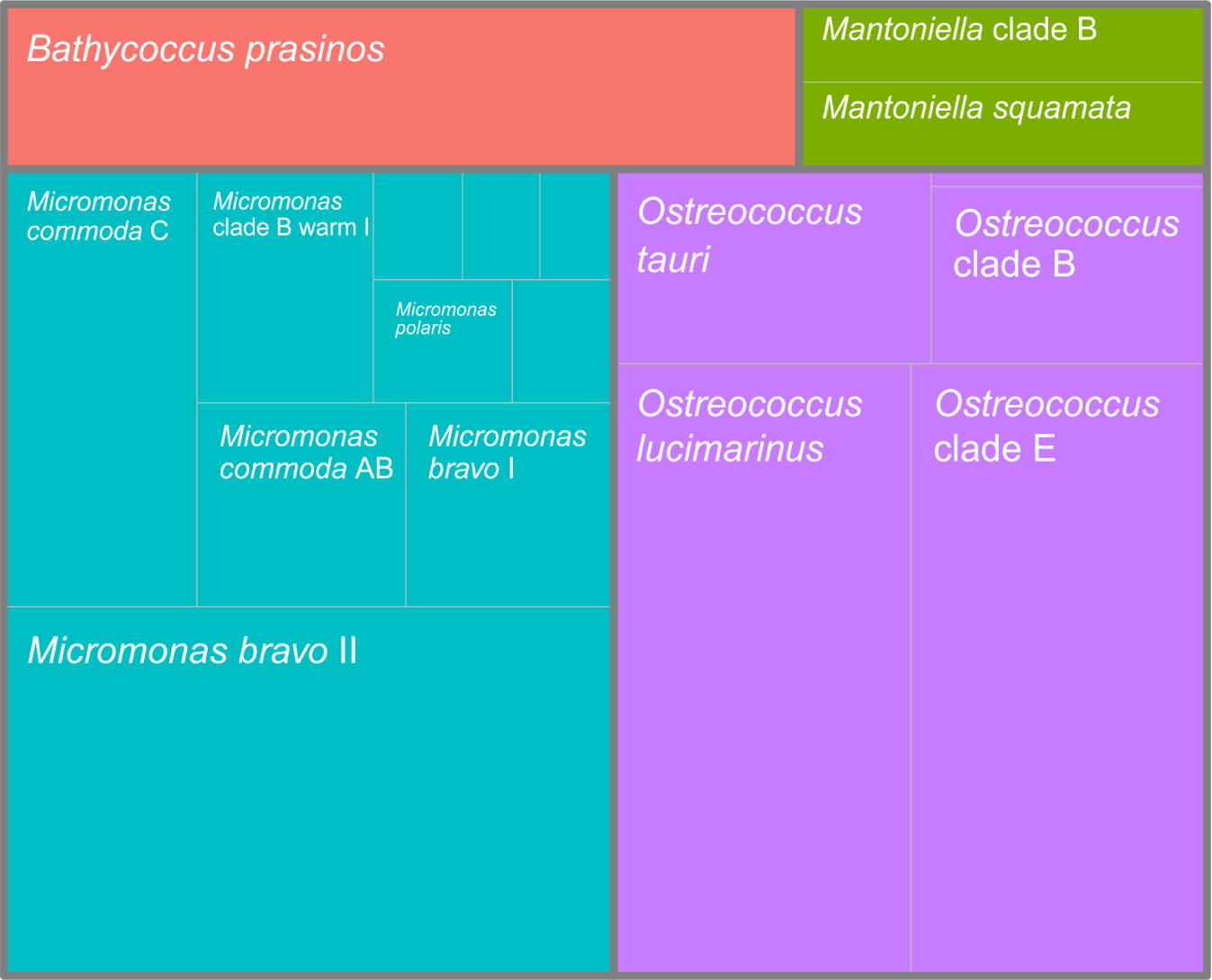
Treemap of the Mamiellophyceae genera and clade contribution for the OSD2014 LGC dataset. Only selected ASVs with more than 200 reads and stations with more than 100 reads were taken into account.

**Table 1.**
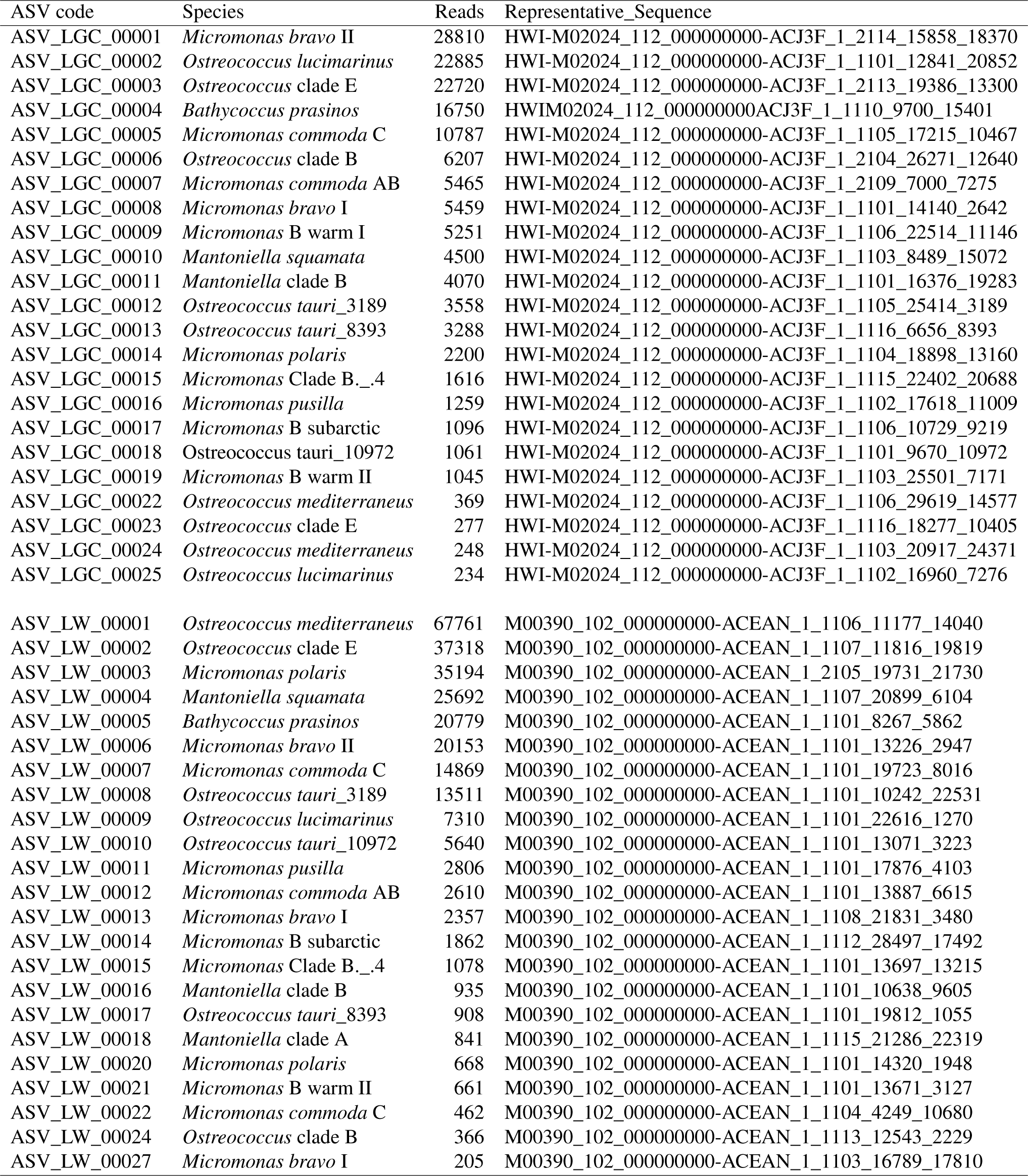
Major Mamiellophyceae ASVs from the LGC and LW datasets: assignation, total abundance and representative sequences name.

### Ostreococcus

4,223 ASVs from the LGC dataset were assigned to the genus *Ostreococcus*, among which 10 corresponded to more than 200 reads (Table 1). For each of these abundant LGC ASVs, we always found a corresponding ASV in the LW dataset. These sequences constituted five clades (Fig. 2A), four already described and a new one that we named clade E following the initial nomenclature of *Ostreococcus* clades^19^. The same tree topology was recovered with ML and Bayesian methods. Alignments (Fig. 2B) confirmed, that the V4 region exhibited clear signatures to delineate the five *Ostreococcus* clades. In this region, the genetic variation between clades (Table S2) is too low to discriminate the different clades when V4 OTUs are built at 99 % identity threshold, except for *O. mediterraneus* (98.3 %). In terms of overall distribution, Ostreococcus was completely absent from high latitudes beyond 60 °N (Fig. 3). *O. “lucimarinus*” was present in the largest number of samples (Fig. S1). In general none of the different Ostreococcus species/clades seemed to co-occur, with the exception of *O. tauri* and *O. “lucimarinus”* and to a lesser extent, clade B and E (Fig. S2).

**Figure 2.**
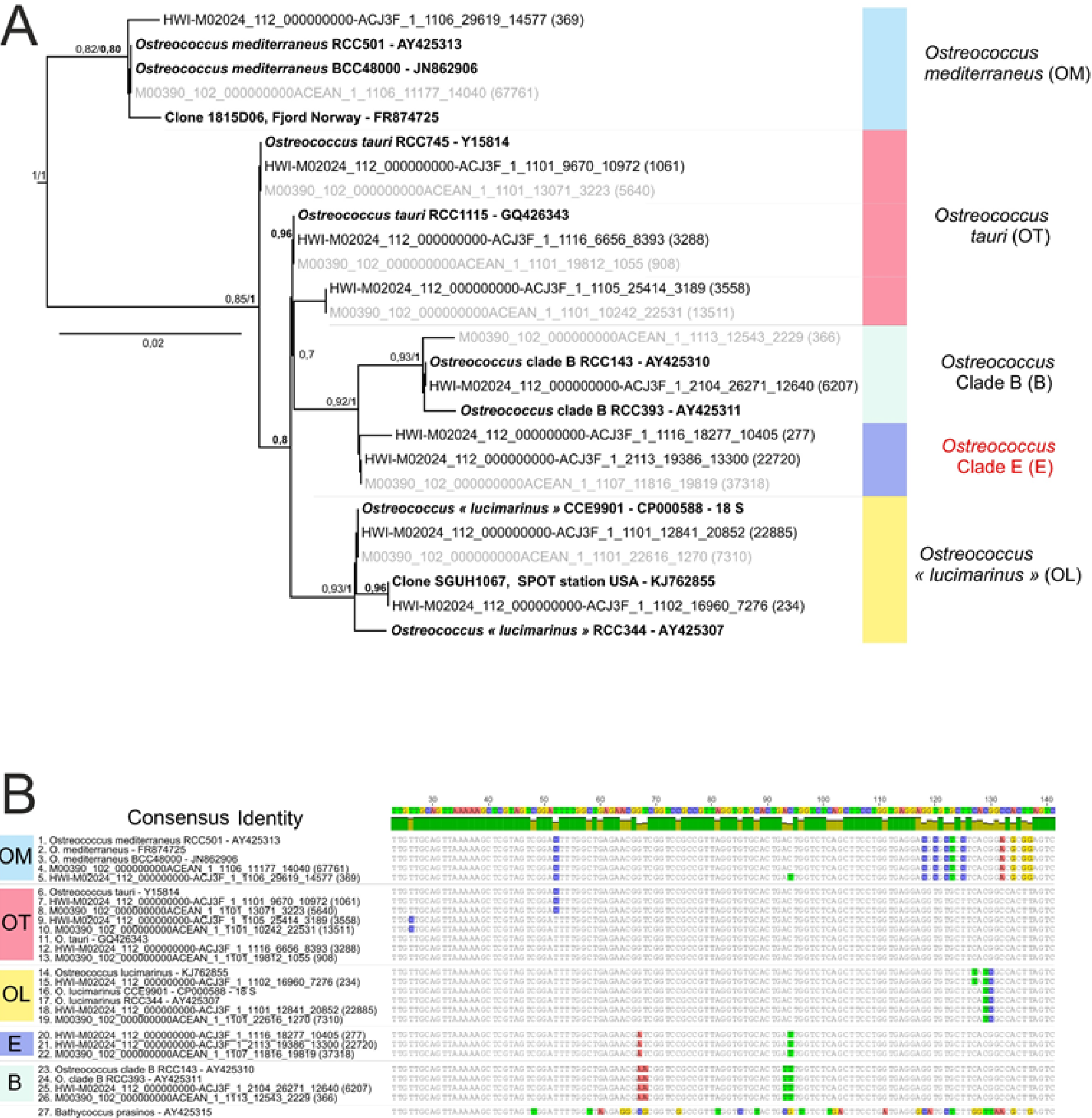
Phylogenetic diversity within the genus *Ostreococcus*. A. Phylogenetic tree of 26 *Ostreococcus* V4 regions of the 18S rRNA gene. The tree was rooted with *B. prasinos* and only ML bootstrap values higher than 70 % are represented and Bayesian posterior probabilities are in bold. Reference sequences from GenBank are in bold, ASVs from the LW dataset (starting with M) are in grey and ASVs from LGC in black (starting with H). Numbers in brackets correspond to the number of reads for each ASV. Only ASVs represented by more than 200 reads were taken into account. B. Alignment of 26 Ostreococcus V4 regions, the alignment is 341 bp long, but only the main signatures are shown (between positions 20 and 140 of the original alignment).

**Figure 3.**
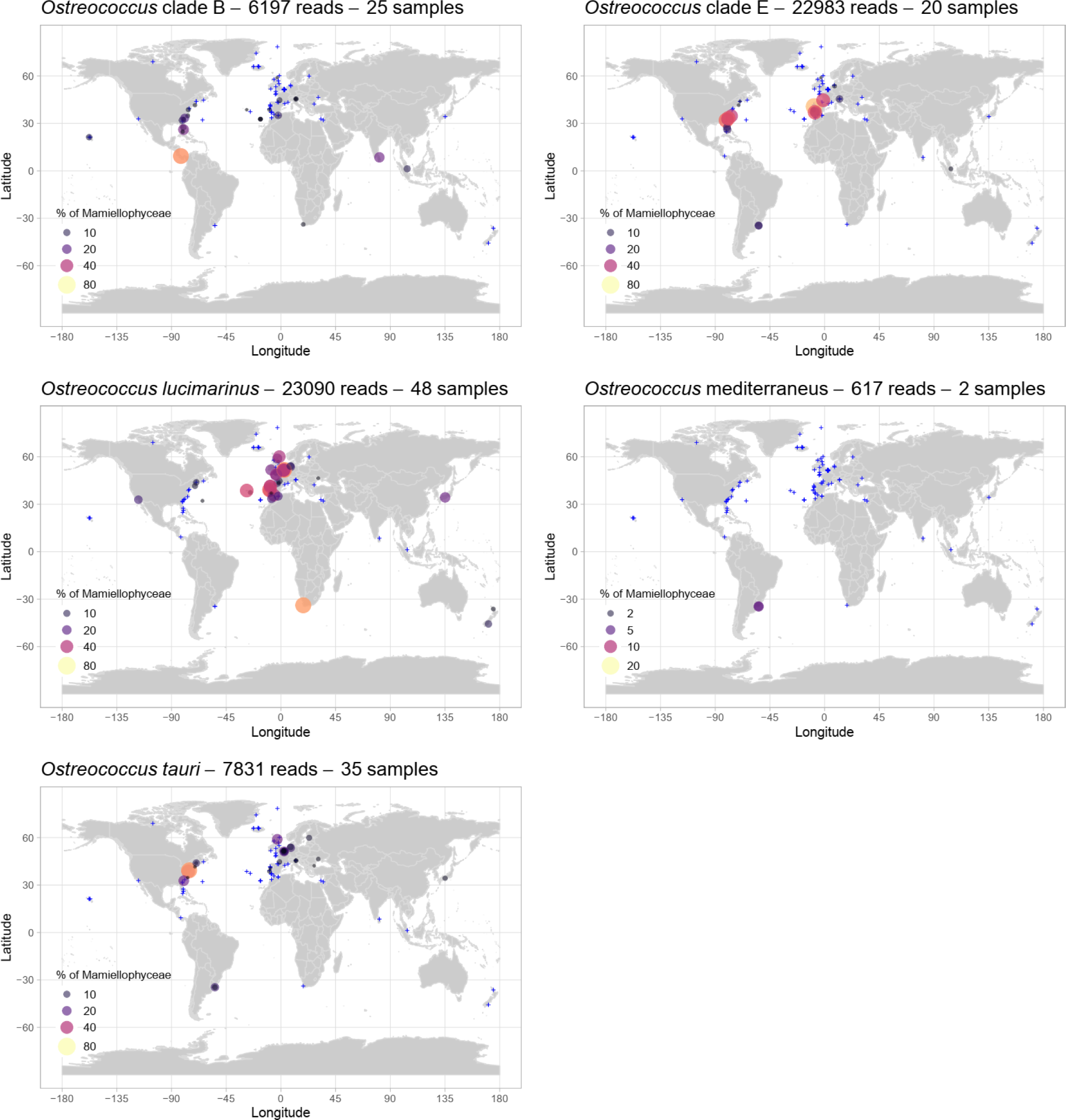
Distribution of the major Ostreococcus ASVs for OSD2014 (LGC). The three major O. tauri ASVs have been pooled together. Circle surface corresponds to the percentage of ASV reads relative to the total number of Mamiellophyceae reads. Samples for which the ASV contribution was lower than 1 % are represented by blue crosses. A zoom on European waters is provided in Fig. S3.

Three ASVs were assigned to the first described species of *Ostreococcus, O. tauri*, with a number of reads ranging from 3,500 and 1,000 (Table 1). These unique sequences did not form a monophyletic clade in either ML and Bayesian trees (Fig. 2A). The two major ASVs (ASV LGC 00012 and ASV LGC 00013) were close to the 18S rRNA sequence recovered from the genome of *O. tauri* (GQ426344). The third ASV (ASV LGC 00018) was identical to sequence Y15814 which was obtained more than 20 years ago for the type strain of *O. tauri*. The V4 alignment (Fig. 2B) reveals two signatures for *O. tauri. O. tauri* ASVs represented more than 1 % of Mamiellophyceae reads at 35 stations and were abundant on the East Coast of the US, in the Baltic Sea, the Adriatic Sea (Venice lagoon), the Black Sea and in Uruguay lagoons (Fig. 3 and S3). Some of these samples correspond to brackish environments (e.g. OSD35 – 8.97 psu, OSD36 – 7.42, OSD39 – 24.3 psu, OSD186 – 7.20 psu) which are similar to the Thau lagoon (highly variable salinity isfrom 24 to 38 psu), where *O. tauri* was initially isolated,^20^. Surprisingly *O. tauri* was absent from the western Mediterranean Sea despite the fact that strains have been isolated from there^21^. The existence of several ASVs suggests that *O. tauri* might be a complex of species, that need to be better distinguished and that could be adapted to different ranges of salinity.

Two major ASVs were assigned to O. *”lucimarinus”*, which was initially described as a high light adapted clade^22^, corresponding to 23,119 reads (Table 1) and representing up to 64 % of the Mamiellophyceae reads off South Africa (Robben Island, OSD133). It dominated Atlantic and North Sea European coastal stations (Belgium OSD183 62 % and 184 44 %, Portugal OSD115 53 %) and represented 40 % of Mamiellophyceae reads at one of the 3 Azores stations (OSD98, Fig. 3). In contrast, *O. “lucimarinus”* was almost totally absent from the Mediterranean Sea and tropical waters (Fig. 3 and S3). This distribution is coherent with observations by qPCR (clade OI according to Demir-Hilton et al. 2011^39^, see below) as a cold mesotrophic coastal clade.

A single ASV was assigned to *Ostreococcus* clade B, initially described as low light adapted clade^22^, representing 6,207 reads and reaching 62 % of Mamiellophyceae reads off Panama (OSD51, Fig. 3). Clade B contribution to Mamiellophyceae was higher than 10 % at 7 tropical and sub-tropical stations from a range of oceans (OSD60 South Carolina, OSD25, 37, 51 Florida, OSD95 Singapore, OSD122 Red Sea, OSD147 Sri Lanka, Fig. 3), which is consistent with previous results obtained by qPCR^39,40^.

The novel *Ostreococcus* clade E was represented by a single ASV in the LGC dataset (22,720 reads) and a similar sequence was also found in the LW dataset, suggesting that it was not an artefact. Clade E sequence is very close to that of clade B, with 2 clear differences (Fig. 2B) and is 100 % similar to a single Genbank sequence (MH008654) obtained also by Illumina sequencing from South China Sea waters. The V4 sequence of clade E is 99.4% similar to that of clade B, such that these two clades may have been lumped together in many metabarcoding studies which do not consider ASVs but OTUs. Clade E could be locally dominant in OSD samples representing up to 70 % of the Mamiellophyceae reads (OSD111, off Portugal). It dominated coastal warm temperate stations (Fig. 3) on both sides of the Atlantic Ocean (Southern USA, OSD39, 58 and 143; Portugal OSD81, 111, 117, 153; France OSD154) and the Mediterranean Sea (Adriatic Sea off Venice, OSD69). It is surprising that no culture has ever been obtained for this new clade but it could be due to the fact that it requires specific conditions to grow.

Finally, two ASVs with a low number of reads (617 in the LGC dataset, Table 1) were assigned to the species *O. mediterraneus*, which was only found in significant proportion in a lagoon of the coast of Uruguay (OSD149 5 % and 150 6 %, Fig. 3). This distribution is coherent with the fact that almost all strains of *O. mediterraneus* have been isolated from coastal lagoons along the Mediterranean Sea coast^21^, suggesting that this species may be restricted to very specific environments with fluctuating salinity.

Two sets of qPCR primers and probes have been previously designed^39^ based on available V4 sequences from strains in culture in order to discriminate two *Ostreococcus* groups (OI and OII). The OI set targets *O. “lucimarinus”* but also recognizes *O. tauri*^39^ which has 2 mismatches (Fig. S4), while the OII group targets clade B. Interestingly the new clade E has four mismatches to set OI and two mismatches to set OII. Some of these mismatches are located on the 3’ end of the forward primer which may prevent either of these sets to recognize *Ostreococcus* clade E, although this would have to be tested once clade E strains become available. If none of these sets recognizes clade E, it would be necessary to design new qPCR sets for clade E. If one or two of these sets recognize clade E, it will cast serious doubts on the validity of previous analyses^25,39,40^ since clade E seems to have a different distribution from both O. “lucimarinus” and clade B.

### Micromonas

4,285 unique OSD LGC sequences were assigned to the genus *Micromonas* with 10 corresponding to more than 200 reads. Phylogenetic analysis and V4 signatures allowed to divide these sequences into 10 major clades (Fig. 4) corresponding to the species, candidate species and clades recently described by Simon et al.^15^ with the exception of a new clade not seen previously, named clade B sub-arctic. *M. commoda, M. bravo* and clade B warm (candidate species 2^15^) could be each divided further into sub-clades: *M. commoda* AB and C^41,42^, *M. bravo* I and II, and B warm I and II. The well supported tree topology was recovered with both ML and Bayesian methods. The genetic divergence between clades is larger than 1 % for almost all the clade pairs (Table S3) and allows to distinguish all clades when using a 99 % identity threshold except for *M. commoda* AB and C (99.2 % identity), *M. polaris* and the new clade B sub-arctic (99.2 %), and *M. commoda* and clade B warm. *M. commoda* C was present in the largest number of samples (Fig. S1). Most species/clades did not co-occur, with the exception of M. bravo I and II (Fig. S2).

**Figure 4.**
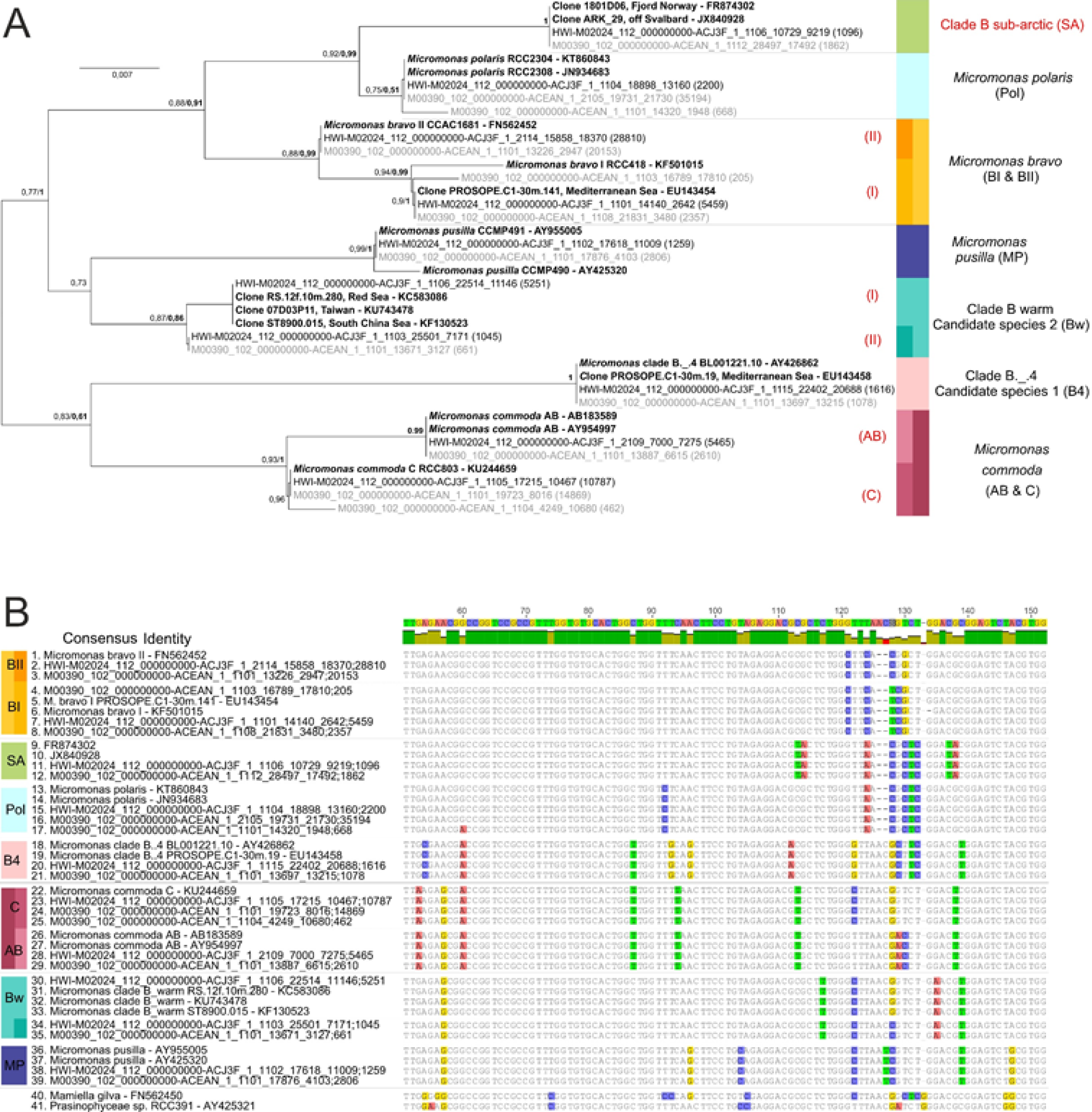
Phylogenetic diversity within the genus *Micromonas*. A. Phylogenetic tree of 39 *Micromonas* V4 regions of the 18S rRNA gene (Fastree). The tree was rooted with Mamiellales (RCC391, AY425321 and Mamiella gilva, FN562450). Legend as Fig. 2. B. Alignment of 39 *Micromonas* V4 regions, the alignment is 327 bp long, but only the main signatures are shown (between positions 50 and 150 of the original alignment).

The major *Micromonas* ASV represented by 28,810 reads, was assigned to *M. bravo* II (Fig. 4). *M. bravo* is a newly described species^15^ which was previously part of the B clade^19^. This ASV represented up to 60 % of the Mamiellophyceae reads (Fig. 5 and S5) in the Black Sea (OSD131) and off Portugal (OSD102). It dominated most of Mediterranean Sea stations, some North European stations and to a lower extent some Pacific stations (off California OSD43 14 % and Hawaii OSD144 28 %). It was the only *Micromonas* ASV that represented more than 10 % of Mamiellophyceae reads off Japan (OSD124 29 %). *M. bravo* I ASV (5,459 reads) represented more than 10 % of Mamiellophyceae at 3 stations along the European coast (in particular 47 % in the English Channel off Plymouth, OSD1).

**Figure 5.**
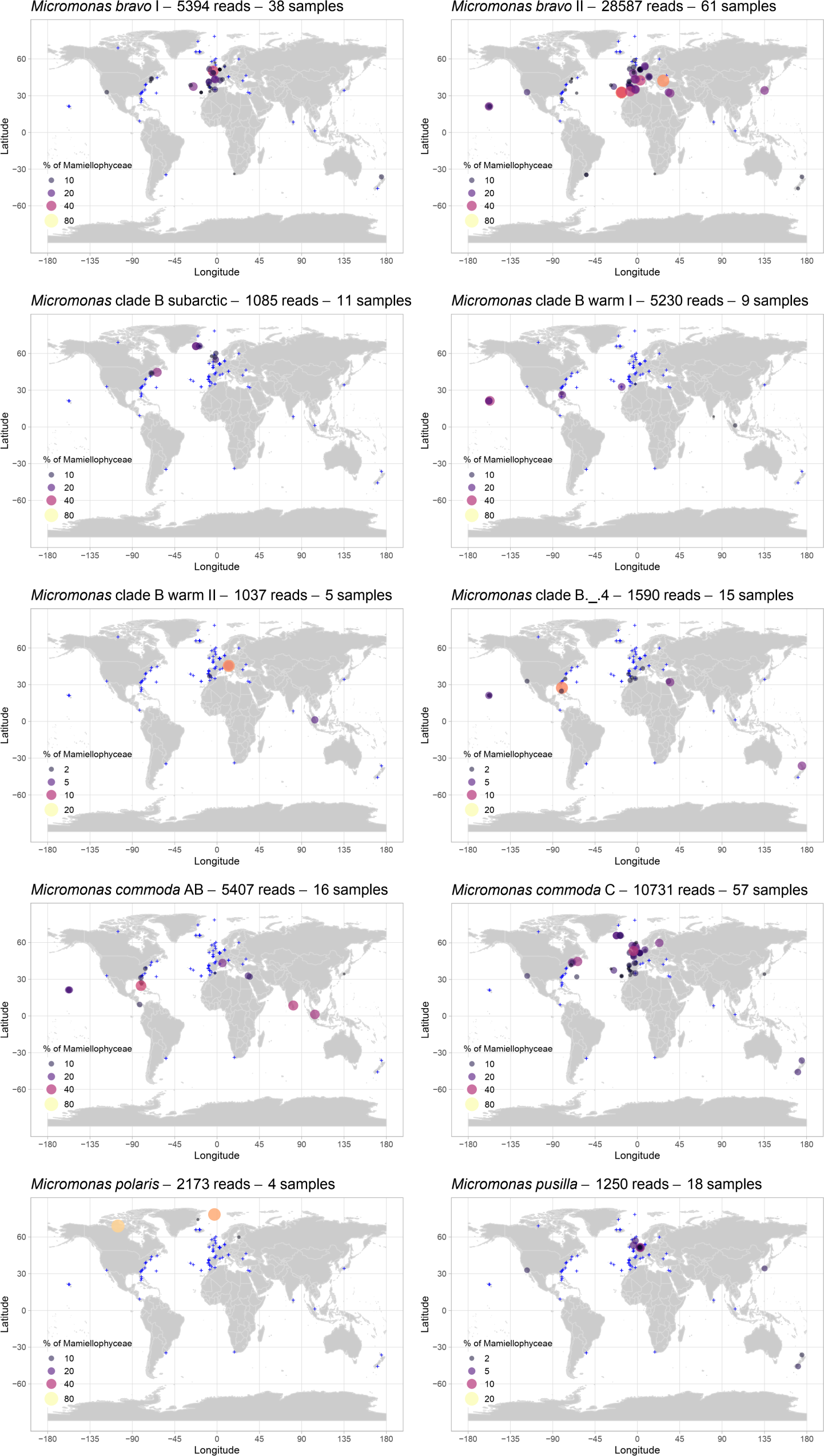
Distribution of the major *Micromonas* ASVs for OSD2014 (LGC). Legend as Fig. 3. A zoom on European waters is provided in Fig. S5.

Two LGC ASVs with a single base pair difference (Fig. 4B) were closely related to the references sequences of the clade B-warm, corresponding to “candidate species 2” from Simon *et al*.^15^ and to “clade VI” from Lin *et al*. 2017^43^. They are referred here as B warm I and B warm II. Clade B warm I ASV (5,251 reads) which matched the original “candidate species 2” sequences (Fig. 4B) contributed to more than 10 % of the Mamiellophyceae reads at 5 stations from tropical or warm waters (Fig. 5: at the 3 Hawaii stations (OSD56 38 %, 57 20 %, 144 18 %), off Florida (OSD37 23 %), off Portugal (OSD101 23 %). It was also found in Singapore (OSD95 5 %). This distribution confirmed the data from available Genbank sequences^15^: one representative strain has been isolated in Mediterranean Sea in summer (http://roscoff-culture-collection.org/rcc-strain-details/1109RCC1109) and environmental clones have been recovered in the Red Sea^44^, South China Sea (“unknown clade”^45^) and off Taiwan (Micromonas clade VI^43^). The second ASV from B warm II did not match any GenBank sequence (Fig. 4B) and was recorded in 1,045 copies, representing more than 1 % of the Mamiellophyceae reads at three stations in Venice lagoon (OSD47, 69, 70 16), off Portugal (OSD81) and in Singapore strait (OSD95).

*M. commoda* C ASV (10,787 reads) represented more than 1% of the Mamiellophyceae at 57 stations (Fig. S1) and up to 35 % off the Atlantic coast of Canada (OSD152). It was found in the North Atlantic up to Iceland and as well off New Zealand (Fig. 5 and S5). In contrast, it was almost completely absent from the Mediterranean Sea as well as from tropical stations. *M. commoda* AB (5,465 reads) was above 1% at a much lower number of stations (16, Fig. 1) and contributed up to 40 % of the Mamiellophyceae off Sri Lanka (OSD147). It was distributed in tropical and subtropical waters (Fig. 5 and S5) in particular off Florida, Singapore and Hawaii as well as in the Eastern Mediterranean Sea off Israel. *M. commoda* was described by Van Baren *et al*.^46^ who just mentioned that this species was not recorded in high latitudes yet (beyond 60 °North and South). The species was then revised by Simon *et al.*^15^, who described the distribution of this species as ubiquitous based on available Genbank sequences. The genetic variability within this species had already been highlighted previously^41,47^ and is comforted by the OSD data since the two clades AB and C have clearly distinct distributions. Simon *et al.*^15^ proposed the hypothesis that speciation events may ongoing within *M. commoda*.

Thz ASV (1,616 reads)corresponding to *Micromonas* clade B. .4 (”candidate species 1” according to Simon *et al.* 2017^15^) reached 15 % off Florida (OSD29) and was found in rather warm waters (off Hawaii, Israel, Morocco, Fig. 5) which matches the distribution of Genbank sequences that had been previously recovered from the Mediterranean and Red Seas as well from the Pacific Ocean^15^.

*M. polaris* ASV (2,200 reads) was the major Mamiellophyceae contributor (Fig. 5) at 2 stations in Arctic waters (73 % in Nunavut OSD105, 66 % in Fram Strait OSD146) and also present in the Gulf of Finland (OSD30). This consistent with the current knowledge on this species. *M. polaris* was first isolated from the Arctic Ocean^48^ and shown to be the dominant pico-eukaryote in the Beaufort Sea in the summer^49^. It has also been recently recorded in the Southern Ocean^50^, although its presence appears less prevalent since it is absent from environmental Antarctic clone libraries^6^. A new *Micromonas* clade genetically close to *M. polaris* (B sub-arctic, 1,096 reads) had maximum contribution off Canada (Bedford Basin OSD 152 32 %) and represented more than 10 % of Mamiellophyceae reads at 4 sub-arctic stations off Maine and Iceland as well as at a temperate location off UK coast in the North Sea (Fig. 5). This ASV is 100 % similar to Genbank sequences recently obtained in the White Sea^51^.

Finally, the ASV corresponding to the first described *Micromonas* species, *M. pusilla*, was found in low abundance (1,259 reads) mostly in temperate locations (Fig. 5) corresponding to the environment from which it was initially described and isolated (e.g. CCMP490 isolated from Woods Hole, USA, and CCMP491 from the English Channel, UK^41^).

### Bathycoccus

The ASV corresponding to *Bathycoccus* was both the most abundant (24,391 reads) as well as the most prevalent, exceeding 1 % of the Mamiellophyceae reads at 72 stations (Fig. S1) distributed all over the coastal ocean from tropical to polar waters (Fig. 6 and S6). This very global distribution of the genus matches what has been observed based on the TARA Oceans dataset^24^ where the metagenome of *Bathycoccus* was recovered at a wide rang of stations. *Bathycoccus* is now known to be composed of two cryptic species with identical 18S rRNA sequences (that therefore cannot be distinguished in the OSD dataset) but differences in the ITS gene as well as at the genomic level^24,52^. The distribution of these two cryptic species (BI-genome RCC1105 and BII-genome TOSAG39-1) determined by metagenomic analysis and qPCR suggest that BI could be coastal, while BII could be adapted to warmer oceanic waters^24,25,53^.

**Figure 6.**
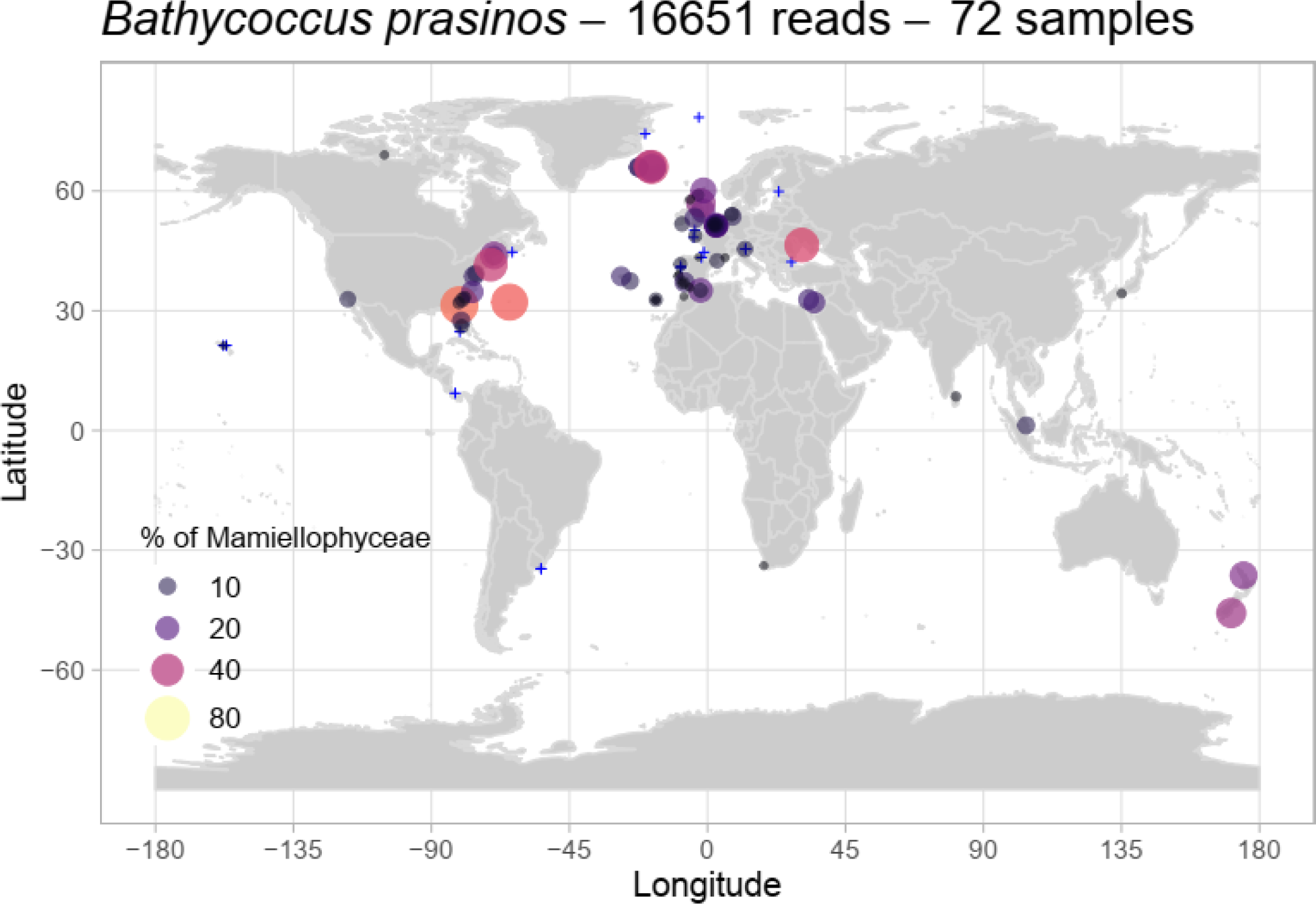
Distribution of the major *Bathycoccus* ASV for OSD2014 (LGC). Legend as Fig. 3. A zoom on European waters is provided in Fig. S6.

### Mantoniella

8,570 reads corresponding to two ASVs with more than 200 reads could be assigned to the genus *Mantoniella*. Besides the morphological species *M. squamata*^3,16^ and *M. antarctica*^18^, no clades based on 18S rRNA gene sequences have been yet described^10,19^. However, the V4 hyper-variable region of the 18S rRNA gene using publicly available reference sequences as well as OSD ASVs highlights two new *Mantoniella* clades that we named A and B, both with well supported phylogenies (Fig. 7A) and with clear sequence signatures (Fig. 7B).

**Figure 7.**
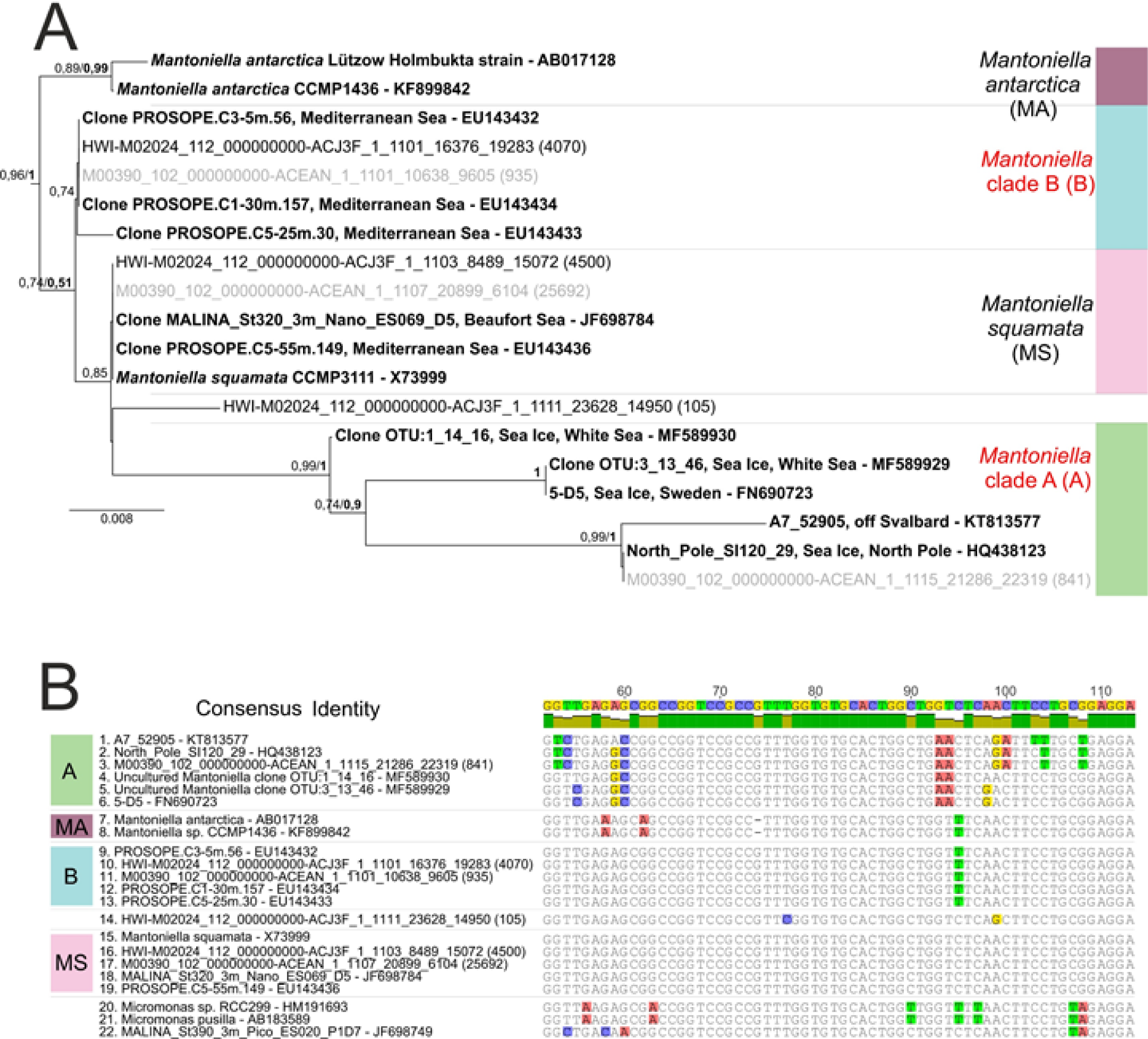
Phylogenetic diversity within the genus *Mantoniella*. A. Phylogenetic tree of 19 Mantoniella V4 regions of the 18S rRNA gene (Fastree). The tree was rooted with the 3 *Micromonas* sequences (AB183589, HM191693, JF698749) and only bootstrap values higher than 70 % are represented. Legend as Fig. 2. B. Alignment of 19 *Mantoniella* V4 regions, the alignment was 368 bp long, but only the main signatures are shown (around the 20th and 141th position of the original alignment).

The most abundant ASV corresponded to the species *M. squamata* (4,500 reads) represented up to 70 % of the Mamiellophyceae reads off Greenland (OSD80, Fig. 8 and S7) while at all other stations it contributed always less than 10 %. However it was found at stations with very different environmental conditions, including off Hawaii and in a lagoon on the coast of Uruguay (Fig. 8). This is consistent with previous descriptions of *M. squamata* as a cosmopolitan species^3^ and several studies reported in particular its presence in Northern high latitudes^49,54^.

**Figure 8.**
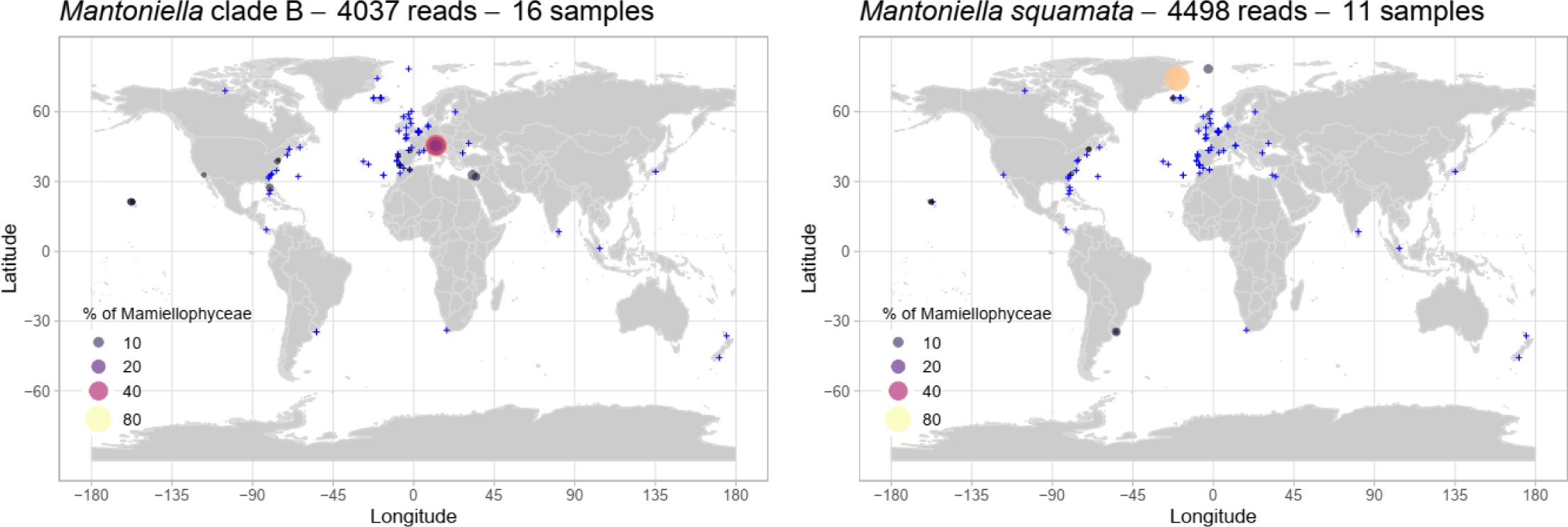
Distribution of the major *Mantoniella* ASVs for OSD2014 (LGC). Legend as Fig. 3. A zoom on European waters is provided in Fig. S7.

The ASV corresponding to *Mantoniella* clade B (4,070 reads) was more widespread, representing more than 1 % of the Mamiellophyceae reads at 16 stations (Fig. S1) especially in the Venice lagoon (OSD47 53 %, OSD70 36 % and OSD69 12 %) but also in other moderately warm waters (Eastern Mediterranean Sea, Gibraltar, California, Hawaii, Fig. 8 and S7), matching Genbank sequences previously found in the Mediterranean Sea^12^.

No ASV corresponding to *Mantoniella* clade A was found in the LGC dataset but one ASV from the LW dataset (841 reads) matched this clade but its only present off Greenland (OSD80) which matched the environment where the other Genbank sequences have been obtained (sea ice from the Arctic Ocean^55^, White Sea^51^ and Baltic Sea^56^) pointing to *Mantoniella* clade A being potentially an ice alga.

Finally no ASV corresponded to the species *M. antarctica* was found either in the LGC or LW dataset. This suggests that this species is probably restricted to Antarctica and was not present at the single station sampled there (OSD187) in the mid of the austral winter where only 85 Mamiellophyceae reads were recorded.

## Limitations of the OSD dataset

The OSD metabarcoding dataset is invaluable to determine the distribution of many phytoplankton groups in coastal waters^9,11,57^. However it must be always emphasized that metabarcoding is not a quantitative method because of biases in the PCR reaction and of the variation in the number of 18S rRNA gene copies per organism^58^ However in the case of Mamiellophyceae, because their size is small, the number of copies is low^58^ and does not vary too much between the different genera (e.g. 2 and 4, copies in *Bathycoccus* and *Ostreococcus*, respectively^59,60^). This supports the use of relative number of reads as a semi-quantitative proxy of Mamiellophyceae contributions. Another potential problem of the OSD dataset is the fact that sampling was done everywhere almost on the same date (around June 21). It has the advantage of providing a snapshot of the world coastal ocean, which allows to work on spatial distribution without the impact of the seasonality, in contrast to oceanographic expeditions such as Tara Oceans^5^ which sampled different ecosystems at different time of the year. One OSD limitation however is that the northern and hemisphere were sampled at opposite sides of the yearly cycle, at the end of the spring and of the fall, respectively. However since sampling was mostly in the Northern hemisphere, this has a low impact on the data interpretation. The existence of phytoplankton cycles driven by temporal changes and nutrient cycles in coastal environment^61–63^ may explain why some species were not found in the OSD dataset in regions where strains or clones corresponding to the same species have been isolated before. As an example, *Bathycoccus* initially isolated from the Gulf of Naples in the Mediterranean Sea^23^ was not recovered at the Naples OSD station (OSD4), where only 2 Mamiellophyceae reads were obtained. Metabarcoding analyses at the Long Term Ecological Research station in the Gulf of Naples show that Mamiellophyceae were absent in June^64^, which explain why they are absent from the OSD dataset. In contrast, analysis of several time series^65^ led to the conclusion that *M. bravo* (previously non arctic B.E.3 clade) dominated the *Micromonas* community in summer and should be adapted to warm well lighted coastal waters which is consistent with what we observed in the OSD dataset for which sampling was done in June.

## Conclusion

Although Mamiellophyceae are clearly the dominant group of green algae in coastal waters^9,11^, previous analysis revealed that their relative abundance could not be related to any environmental variable^9^. Even analysis at the genus level is not sufficient to detect bio-geographical patterns. For example, the genus *Micromonas* is found at virtually all OSD stations (Fig. 5). The only genus which is not ubiquitous is *Ostreococcus* which is not found in polar regions. In contrast, analysis at the species/clade levels allows detecting some very clear patterns in particular with respect to latitudinal distribution (Table 2. Some species/clade have quite restricted latitudinal ranges, e.g. *M. polaris* only found in Arctic samples, *M. pusilla* only found in temperate waters or *Micromonas* clade B warm I only in tropical waters. Others are much more wide spread, for example *Ostreococcus* clade B which extends from temperate to tropical waters. Some taxa seem to be restricted to specific habitats, in particular *O. mediterraneus* to coastal lagoons. The case of the Mediterranean Sea is also interesting. It has been previously shown to harbour specific taxonomic groups such as Chlorodendrophyceae^9^, clade A6 of Chloropicophyceae^11^ or the Bolidophyceae *Triparma mediterranea*^66^. Here, we found that some temperate species such as *M. pusilla* or *Ostreococcus* clade E were completely absent from this region but we did not find any species/clade restricted to it.

**Table 2.**
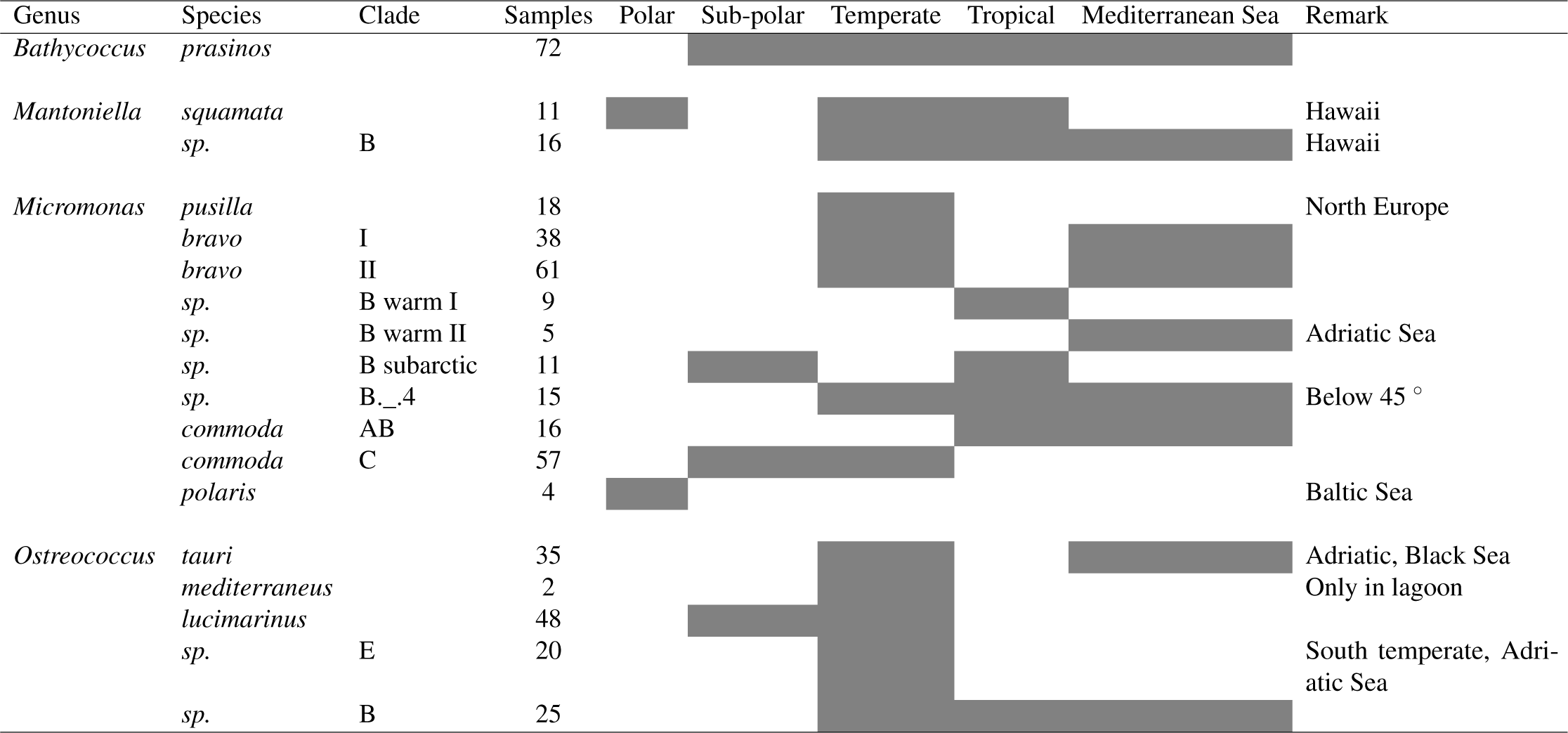
Summary of the coastal distribution of Mamiellophyceae species and clades. The column indicate the number of samples where the species/clade represented more than 1% of Mamiellophyceae

Genetic diversity is quite different between the four genera that we examined. While *Bathycoccus* is composed of a single clade at least based on the 18S rRNA gene, while *Micromonas* seems extremely diversified with a large number of clades. Although most of the clades found in OSD have been observed before, we uncovered some new diversity. One species, *M. bravo* was split into two sub-clades, I and II, that seem to have distinct distribution. Two new clades were uncovered. *Micromonas* B subarctic seems to be restricted to a specific latitudinal band (roughly 45 to 65 °N). *Ostreococcus* clade E seems also very interesting since it is very abundant and has a distinct distribution from the closely related *Ostreococcus* clade B by being restricted to temperate waters while clade B is found tropical waters. Since some oceanic regions have not been covered by the OSD dataset, it is possible that yet undiscovered Mamiellophyceae clade/species may exist in particular in the Southern hemisphere. In order to obtain more information on the new clades reported here and to determine whether they correspond to potentially novel species, several strategies are possible. First, now that their geographical distribution begins to be known, we may target specific environments and obtain isolates. This will allow analyzing finer resolution markers such as the ITS^15^ and determine physiological preferences, for example in terms of temperature^67^. Another strategy for clades that are hard to isolate in culture would be to determine longer sequences of the ribosomal operon including in particular the ITS region directly from the environment either by long amplicon PCR using novel sequencing technologies such as Nanopore^68^ or by extracting them from existing metagenomics datasets such as those obtained during the Tara Oceans project^24^.

## Acknowledgements

Financial support for this work was provided by the European Union projects MicroB3 (UE-contract-287589), the ANR PhytoPol (ANR-15-CE02-0007) and TaxMArc (Research Council of Norway, 268286/E40). MT was supported by a PhD fellowship from the Université Pierre et Marie Curie and the Région Bretagne (ARED GreenPhy). We would like to thank the Ocean Sampling Day consortium for providing sequence data and the ABIMS platform in Roscoff for access to bioinformatics resources.

## Author contributions statement

D.V. and M.T. conceived the study. D.V. produced some of the figures and edited the paper. M.T. analyzed the data, produced some of the figures and wrote the initial draft of the paper. All authors reviewed the manuscript.

## Additional information

### Data availability

Scripts for mothur, Mamiellophyceae OTU sequences, alignments, assignation and abundance for the LGC and LW datasets are provided as supplementary files on Figshare: https://figshare.com/s/6e074ed2 R processing script is detailed at https://vaulot.github.io/papers/OSDMamiello.html and provided at https://github.com/vaulot/Paper-2018TraginMamiellophyceae-R-scripts

